# Phenomic selection: a low-cost and high-throughput method based on indirect predictions. Proof of concept on wheat and poplar

**DOI:** 10.1101/302117

**Authors:** Renaud Rincent, Jean-Paul Charpentier, Patricia Faivre-Rampant, Etienne Paux, Jacques Le Gouis, Catherine Bastien, Vincent Segura

## Abstract

Genomic selection - the prediction of breeding values using DNA polymorphisms - is a disruptive method that has widely been adopted by animal and plant breeders to increase productivity. It was recently shown that other sources of molecular variations such as those resulting from transcripts or metabolites could be used to accurately predict complex traits. These endophenotypes have the advantage of capturing the expressed genotypes and consequently the complex regulatory networks that occur in the different layers between the genome and the phenotype. However, obtaining such omics data at very large scales, such as those typically experienced in breeding, remains challenging. As an alternative, we proposed using near-infrared spectroscopy (NIRS) as a high-throughput, low cost and non-destructive tool to indirectly capture endophenotypic variants and compute relationship matrices for predicting complex traits and coined this new approach “phenomic selection” (PS). We tested PS on two species of economic interest (*Triticum aestivum* L. and *Populus nigra* L.) using NIRS on various tissues (grains, leaves, wood). We showed that one could reach predictions as accurate as with molecular markers, for developmental, tolerance and productivity traits, even in environments radically different from the one in which NIRS were collected. Our work constitutes a proof of concept and provides new perspectives for the breeding community, as PS is theoretically applicable to any organism at low cost and does not require any molecular information.

**ARTICLE SUMMARY:** Despite its widely adopted interest in breeding, genomic selection - the prediction of breeding values using DNA polymorphisms - remains difficult to implement for many species because of genotyping costs. As an alternative or complement depending on the context, we propose “phenomic selection” (PS) as the use of low-cost and high-throughput phenotypic records to reconstruct similarities between genotypes and predict their performances. As a proof of concept of PS, we made use of near infrared spectroscopy applied to different tissues in poplar and wheat to predict various key traits and showed that PS could reach predictions as accurate as with molecular markers.

## INTRODUCTION

To meet the world’s current and future challenges, especially in terms of food and energy supplies, there is a great need to develop efficient crop varieties, livestock races or forest materials through breeding. Until recently, the selection of promising individuals in animal and plant breeding was mostly based on their phenotypic records. This approach was a strong limit to genetic progress as the high costs of phenotyping strongly constrain the number of candidates that can be evaluated, especially when there are interactions between individuals and environments that necessitate the evaluation of selection candidates in various environments. Another strong constraint – typical in perennial crops, trees or animals – is that it can sometimes take several years to evaluate phenotypes, which increases the duration of selection cycles. These limitations are some of the main reasons why genomic selection (GS) has become so popular in the last two decades. Its principle is based on a combination of phenotypic records and genome-wide molecular markers to train a prediction model that can in turn be used to predict the performances of – potentially unphenotyped – individuals (Meuwiseen *et al.* 2001). We can thus select more individuals faster, which increases selection efficiency. The development of high-throughput genotyping tools at decreasing costs has made GS possible for many animal and plant species. It can be used both in pre-breeding to screen diversity material (Crossa *et al.* 2016; Yu *et al.* 2016) and in breeding to make the schemes more efficient (Heffner *et al.* 2010; Meuwissen *et al.* 2013). However, a great number of species are still orphans of any genotyping tool, and for many others, genotyping costs remain a limit to the implementation of GS in pre-breeding and breeding. In addition, genotyping thousands to millions of individuals (potentially each year) is a challenge that consequently remains inaccessible for most species.

One of the reference GS models is the ridge regression BLUP (RR-BLUP, Whittaker *et al.* 2000; Meuwiseen *et al.* 2001) in which a penalized regression is made on all markers simultaneously. This model assumes that the genes affecting the trait of interest are spread across the whole genome and that all of these genes have small effects. Despite its simplicity, this model has been proven one of the most effective in many situations, except for when major genes contribute to trait architecture. Interestingly, this model is equivalent to the genomic BLUP model (G-BLUP, Habier *et al.* 2007; Goddard *et al.* 2009; Hayes *et al.* 2009; Zhong *et al.* 2009) in which markers are used to estimate a realized genomic relationship matrix between individuals, also called kinship. This framework means that we can compress genome-wide information from numerous molecular markers into summary statistics (kinship coefficients between individuals) without diminishing prediction accuracy. Considering this fact, we should ask the question: are there more efficient alternatives than genotyping to estimate the kinship matrix? In the last years, it was proposed to use endophenotypes (MacKay *et al.* 2009) such as transcripts (Fu *et al.* 2012; Guo *et al.* 2016; Zenke-Philippi *et al.* 2016; Westhues *et al.* 2017), small RNAs (Seifert et al. 2018) or metabolites (Riedelsheimer *et al.* 2012; Feher *et al.* 2014; Ward *et al.* 2015; Fernandez *et al.* 2016; Xu *et al.* 2016; Guo *et al.* 2016; Schrag *et al.* 2018) as regressors or to estimate kinship. These endophenotypes correspond to different molecular layers between the genome and the phenotype, which permits the integration of interactions and regulatory networks when getting closer to the phenotypes. These kinds of variables have proven to be efficient to predict integrative traits using the same statistical models as those classically used in GS. These regressors have the advantage of capturing expressed genotypes, but they remain too expensive to be routinely applied on the large scales typically dealt with by breeders. It is interesting to note that even with a small portion of the transcripts or metabolites sampled on a single tissue in a single environment and sometimes at very early stages, it was possible to compute kinship matrices allowing to reach predictive abilities similar to those obtained with molecular markers (Riedelsheimer *et al.* 2012; Xu *et al.* 2016). One could thus consider the possibility of using cheaper and easier techniques to capture endophenotypic variations.

Near-infrared spectroscopy (NIRS) is a high-throughput, non-destructive and low-cost method routinely used to estimate reflectance of a sample for numerous wavelengths. This reflectance is mainly related to the presence of chemical bonds in the analysed tissue and as a result is expected to be related to endophenotypes. We suppose that the reflectance at each of the numerous wavelengths can be considered as an integration of numerous endophenotypic variations. We thus propose to evaluate the efficiency of NIRS to make predictions with G-BLUP (or equivalently, RR-BLUP) using these traits instead of molecular markers. Numerous studies have demonstrated the usefulness of NIRS for barcoding samples and discriminating species or varieties (Bertrand *et al.* 1985; Adedipe *et al.* 2008; Espinoza *et al.* 2012; Fischnaller *et al.* 2012; Abasolo *et al.* 2013; O’Reilly-Wapstra *et al.* 2013; Meder *et al.* 2014; Lang *et al.* 2017) and have thus suggested that NIRS could be considered as a genetic marker (Cruickshank and Munck 2011) Moreover, some studies have shown that NIRS can capture some genetic variability by estimating the heritability of the spectrum and even mapping corresponding quantitative trait loci (QTL, O’Reilly-Wapstra *et al.* 2013; Posada *et al.* 2008; Diepeveen *et al.* 2012; Hein and Chaix 2014). However, to the best of our knowledge, no studies have proposed using NIRS to perform “phenomic selection” (PS), which we define as the use of high-throughput phenotyping to obtain numerous variables which can be used as regressors or to estimate kinship in the statistical models classically used in GS. We emphasize that the concept of phenomic selection is radically different from the classical use of NIRS prediction. In the classical methodology, NIRS is collected on a sample to make prediction on that particular sample for traits of various complexity (from chemical composition (Foley *et al.* 1998) to yield (Ferrio *et al.* 2005; Cabrera-Bosquet *et al.* 2012; Weber *et al.* 2012; Aguate *et al.* 2017) using a formula that has previously been calibrated. On the other hand in PS, NIR reflectances are considered in the same way as genomic or endophenotypic regressors, at the genotypic level rather than the individual level, which allows making predictions in any environment without having any environment specific NIRS. In PS we suppose that once NIR reflectances are analyzed in one experiment (collections of seed, a nursery, a trial or a controlled experiment) they could be used as regressors or to estimate a kinship matrix to make predictions in any other experiment, as long as relevant phenotypic data are available to calibrate the statistical model, like in GS with molecular markers.

There are several advantages to this approach. One can obtain NIRS for any plant or animal species at a lower cost than genotyping and potentially without particular treatment of the samples prior to the analysis such as DNA or RNA extraction. One can also obtain NIRS directly in the field thanks to portable devices (Ecarnot *et al.* 2013; Teixera dos Santos *et al.* 2013) or autonomous high-throughput vectors, such as phénomobiles (Madec *et al.* 2017) that generate hyperspectral images (Diago *et al.* 2013; Peerbhay *et al.* 2013). NIRS can even be obtained non destructively on seeds before sowing. As a result, prediction-based selection would be possible for any species and at a low enough cost to make it interesting to implement, even if its results are less accurate than those of GS. As a proof of concept of PS, we report an evaluation of the usefulness of NIRS for predicting quantitative traits of economic interest within two different species, a tree (poplar) and a cereal (winter wheat) using various tissues (grains, leaves, wood) and under different environments, and compare the results to those of a GS prediction based on several thousand SNPs.

## MATERIALS AND METHODS

### Data

#### Genetic material and experimental designs

*Wheat:* The panel was composed of 228 European elite varieties of winter wheat released between 1977 and 2012, 89% of which have been released since 2000. 72.8% of these varieties are in the panel introduced in Ly *et al.* (2018). The full panel was sown in one trial in Clermont-Ferrand (France) in 2015/2016. This trial was an augmented design with two treatments: one drought treatment under rain-out shelters (DRY), and one irrigated treatment (IRR) next to it. There was a difference of 223 mm in water supply (rainfall and irrigation) between the two treatments at the end of the experiment. For both treatments, the panel was divided into eight blocks of precocity with one replicate within the same block for 64 varieties and no replicates for the other 164, except for four checks, which were replicated three times in each block. Phenotypes and NIRS were collected in these two reference environments. A subset of 161 varieties (together with 59 additional varieties that were not used in the present study because they were not in the panel of 228) were sown and phenotyped in six independent environments located in Estrées-Mons (France, 2011/2012 and 2012/2013) and Clermont-Ferrand (France, 2012/2013) with two treatments corresponding to two levels of nitrogen input (intermediate and high). This subpanel was divided into six groups of earliness and each group was repeated in two blocks. Four checks were present in each block.

*Poplar:* The population was an association population comprising 1,160 cloned genotypes representative of the natural range of the species in Western Europe and previously described (Guet *et al.* 2015; Faivre-Rampant *et al.* 2016; Gebreselassie *et al.* 2017). Clonally replicated trials of subsets of this association population were established in 2008 at two contrasting sites in central France (Orléans, ORL) and Northern Italy (Savigliano, SAV). At each site, a randomized complete block design was used with a single tree per block and six replicates per genotype. Growth data collected in each design clearly indicated that the Italian site was more favorable than the French site (Guet *et al.* 2015; Gebreselassie *et al.* 2017).

#### NIRS data

*Wheat:* NIRS data were obtained on flag leaves and harvested grains from the two treatments of the drought trial in Clermont-Ferrand (France) in 2015/2016. For each variety in each treatment, twenty flag leaves were sampled on one plot at 200 degree days after flowering. The samples were oven dried at 60°C for 48 h. Leaves were milled (Falaise miller, SARL Falaise, France), and the powder was analyzed with a FOSS NIRS 6500 (FOSS NIRSystems, Silver Spring, MD) and its corresponding softwares (ISIscan^TM^ and WINisi^TM^ 4.20). For each variety in each treatment, 200 g of grains harvested at one plot were analyzed with a FOSS NIRS XDS (FOSS NIRSystems, Silver Spring, MD) and its corresponding softwares (ISIscan^TM^ and WINisi^TM^ 4.20). For leaf powder and grain, absorbance was measured from 400 to 2500 nm with a step of 2 nm. 5 varieties were removed from the dataset because their leaf absorbance was abnormal, resulting in a final panel of 223 varieties. The resulting spectra were loaded into R software (R core team, 2017) to be pretreated using custom R code. They were normalized (centered and scaled) and their first derivative was computed using a Savitzky-Golay filter (Savitzky and Golay, 1964) with a window size of 37 data points (74 nm) implemented in the R package signal (signal developers, 2013). In the end, each variety in each treatment was characterized by a transformed spectrum of flag leaf powder and a transformed spectrum of grains.

*Poplar:* NIRS was carried out on wood from stem sections collected at 1 m above ground on 2-year-old trees for 1,081 genotypes in three blocks at Orléans (total of 2,860 samples) and 792 genotypes in three blocks at Savigliano (total of 2,254 samples). After harvest, the wood samples were oven dried at 30°C for several days, cut into small pieces with a big cutter and milled using a Retsch SM2000 cutting mill (Retsch, Haan, Germany) to pass through a 1-mm sieve. The wood samples were not debarked prior to milling. After stabilization, wood powders were placed into quartz cups for NIR collection with a Spectrum 400 spectrometer (Perkin Elmer, Waltham, MA, USA) and its corresponding software (Spectrum^TM^ 6.3.5). For each sample, the measurement consisted of an average of 64 scans done while rotating the cups over the 10,000 cm^−1^ −4,000 cm^−1^ range with a resolution of 8 cm^−1^ and a zero-filling factor of 4, resulting in absorbance data every 2 cm^−1^. The resulting spectra were loaded into R software (R core team, 2017) to be processed using custom R code. They were first restricted to the 8000 cm^−1^ −4000 cm^−1^ range because the most distant part of the spectra (8000 cm^−1^ −10,000 cm^−1^) appeared to be quite noisy. Then, the restricted spectra were normalized (centered and scaled), and their first derivative was computed using a Savitzky-Golay filter (Savitzky and Golay, 1964) with a window size of 37 data points (74 cm^−1^) implemented in the R package signal (signal developers, 2013). Finally, these normalized and derived spectra were averaged by genotype at each site.

#### SNP data

*Wheat:* The 228 wheat varieties were genotyped with the TaBW280 K high-throughput genotyping array described in Rimbert *et al.* (2018). This array was designed to cover both genic and intergenic regions of the three subgenomes. Markers with a minor allele frequency below 1%, or with a heterozygosity or missing rate above 5% were removed. Redundant markers were filtered out. Eventually, we obtained 84,259 SNPs, either polymorphic high resolution or off-target variants, with an average missing data rate of 0.83%. Missing values were imputed as the marker frequency.

*Poplar:* The poplar association population was genotyped with an Illumina Infinium BeadChip array (Faivre-Rampant *et al.* 2016) yielding 7,918 SNPs for 858 genotypes. Missing values were rare (0.35%) and they were imputed with FImpute (Sargolzaei *et al.* 2014). The data were restricted to the subset of 562 genotypes with SNP data and NIRS data at both sites. Within this set, SNPs with a minor allele frequency below 1% were discarded, yielding a final SNP dataset of 7,808 SNPs.

#### Phenotypic data

*Wheat:* The 228 wheat varieties were phenotyped for heading date (HD) and grain yield (GY) at the two environments in which the NIRS analysis was conducted (drought experiment in Clermont-Ferrand 2015/2016). The subpanel of 161 varieties was phenotyped for the same traits in six independent environments. In each environment, the phenotypic data were adjusted for micro-environmental effects using the random effect block and when necessary by modeling spatial trends using two-dimensional penalized spline (P-spline) models as implemented in the R package SpATS (Rodríguez-Álvarez *et al.* 2016).

*Poplar:* The poplar association population was evaluated at each of the two sites for the following traits on up to six replicates by genotype: height at 2 years at Orléans (HT-ORL), circumference at 1 m above ground at 2 years at both sites (CIRC-ORL and CIRC-SAV), bud flush at both sites (BF-ORL and BF-SAV) and bud set at both sites (BS-ORL and BS-SAV) as discrete scores for a given day of the year (see Dillen *et al.* (2009) and Rohde *et al.* (2011) for details on the scales used) and resistance to rust at Orléans (RUST-ORL) as a discrete score of susceptibility on the most affected leaf of the tree and on a 1 to 8 scale. Within each site, the phenotypic data were adjusted for micro-environmental effects using random effect block and/or spatial position when needed following a visual inspection of spatial effects with a variogram as implemented in the R package breedR (Muñoz and Sanchez 2017). Finally, the adjusted phenotypes were restricted to the subset of 562 genotypes with SNP and NIRS data for computing an averaged genotypic value for each trait by genotype within each site for further analyses.

### Genetic variance captured by NIRS

#### Genomic heritability and partition of phenotypic variance along spectra

The estimation of genomic heritability was based on the following bivariate statistical model across environments:

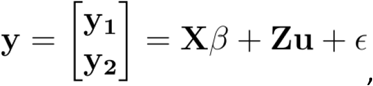

where *y*_1_,*y*_2_ are the phenotypic values (absorbance for a given wavelength) in each environment, *β* is a fixed effect of the environment, **u** is a vector of random polygenic effect with 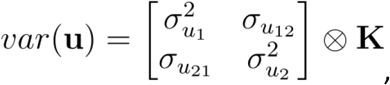 *K* being the scaled realized relationship matrix (see below), *ε* is a vector of independent and normally distributed residuals with 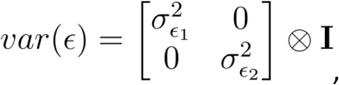 and **X** and **Z** are design matrices relating observations to the effects.

SNPs were used to estimate the genomic relationship matrix (**A**) between individuals, following the formula of VanRaden (Vanraden 2008):

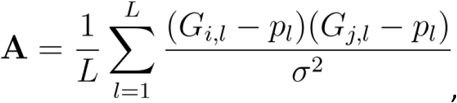

where *G_j,l_* and *G_j,l_* are the genotypes of individuals *i* and *j* at marker *l* (*G*.,*l* = 0 or 1 for homozygotes, 0.5 for heterozygotes), *pl* is the frequency of the allele coded 1 for the marker l, and *σ*^2^ is the average empirical marker variance. *K* was obtained by scaling *A* to have a sample variance of 1 (Kang *et al.* 2010; Forni *et al.* 2011).

Genomic heritability was estimated for each wavelength within each environment (*m*) as follows: 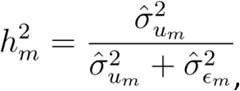 with 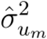 and 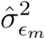 the REML estimates of 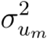 and 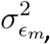 obtained with the Newton-Raphson algorithm implemented in the R package sommer (Covarrubias *et al.* 2016).

Following Yamada *et al.* (1988), the variance/covariance estimates from the previously defined bivariate mixed-model were used to compute estimates of genetic 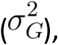 genetic by environment 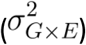 and residual 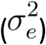 variances across sites as follows:

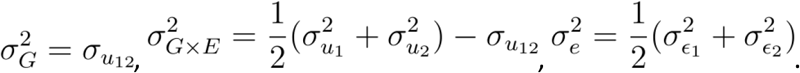

### Association mapping of NIRS reflectance

Association mapping was carried out along spectra considering the absorbance at a given wavelength as a trait in a bivariate setting and using previous estimates of genetic and residual variances (EMMAX philosophy as previously proposed in the multi-trait mixed-model approach (Korte *et al.* 2012)).

### Genomic and phenomic predictions

The efficiency of genomic and phenomic predictions was evaluated by cross-validations in two types of scenarios (Fig. 1). In scenario S1, NIRS analysis and cross-validation were applied to the same environment (Fig. 1 a). In scenario S2, cross-validation was applied to independent environments: the environment(s) in which NIRS was collected and the environment in which the cross-validation was applied (calibration and prediction) were different (Fig. 1 b). In S1, the objective was to limit expensive or labor-demanding phenotyping to a calibration set of reduced size and to predict the remaining individuals using NIRS. In scenario S2, one experiment (or a nursery) was dedicated to collecting the NIRS of the calibration set and the predicted set, and a multi-environment trial was dedicated to phenotyping the calibration set. The main difference between S1 and S2 was that in S1, NIRS could potentially capture both genetic and *G* × *E* variances, leading to environment-specific predictions.

**Figure 1.**
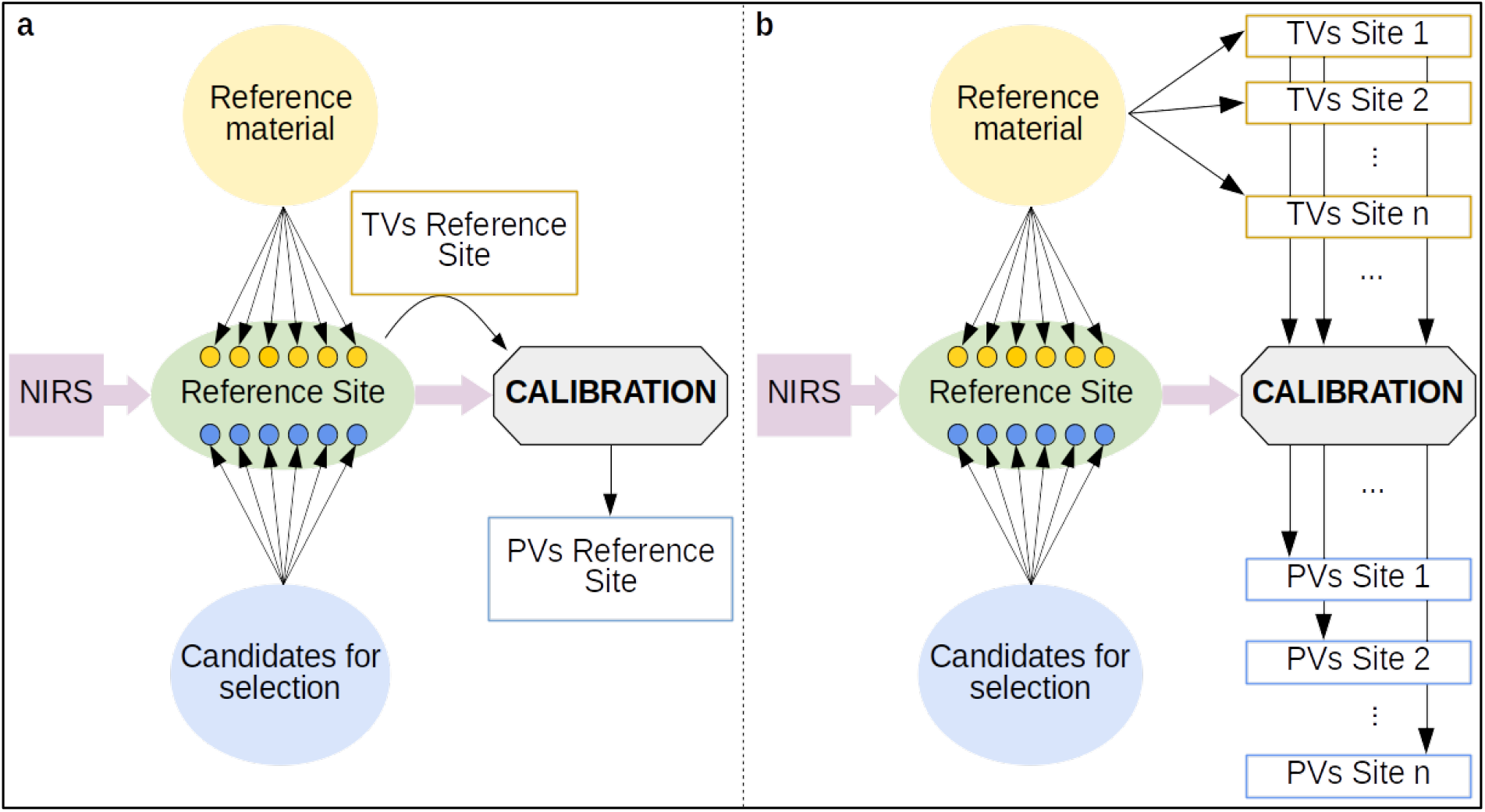
Schematic representation of the concept of phenomic selection, including the two scenarios tested in the present work: (a) S1, where the calibration model is trained with true values (TVs) and NIRS data collected at the same - reference - site and (b) S2, where the calibration model is trained with NIRS data collected at the reference site and TVs from other(s) environment(s). In both scenarios, the outcome of the prediction consists of predicted values (PVs).

For both scenarios, 5- and 8-fold cross-validation procedures repeated 20 times were used for poplar and wheat, respectively. A larger fold-number was considered for wheat in comparison to poplar because the sample size in the wheat dataset (n = 223 in the panel, and n = 161 in the subpanel) was lower than the sample size in the poplar dataset (n = 562). Predictive ability was computed as the Pearson correlation between the predictions and adjusted means. For genomic predictions, we tested two complementary reference models: G-BLUP and Bayesian LASSO (Park and Casella 2008; de los Campos *et al.* 2009. The underlying assumptions of these two models are that the SNP effects are normally distributed for G-BLUP, whereas Bayesian LASSO allows for departure from normality (*i.e.*, SNPs with bigger effects). G-BLUP and Bayesian LASSO were run with the R packages rrBLUP (Endelman 2011) and BGLR (de los Campos *et al.* 2012), respectively. For Bayesian LASSO, the chain was composed of 30,000 iterations with a burn-in of 5,000 iterations, and the hyperparameter λ was chosen as recommended in Table 1 of de los Campos *et al.* (2012). For phenomic predictions, we used RR-BLUP but considered NIRS data instead of molecular markers. Prior to the analysis, the pretreated NIRS matrices were centered and scaled. The shrinkage parameter was estimated within the cross-validation scheme on the calibration set only to avoid overfitting. In other words, the reported predictive abilities for NIRS prediction were unlikely to be overestimated because of model optimization.

### Expected genetic gain with genomic and phenomic selection in a simple example

We ran simulations to illustrate the expected genetic gain with GS and PS that would be achieved in one cycle of selection for various combinations of costs and reliabilities. Reliability was defined as the squared correlation between true breeding values (TBV) and the genomic or NIRS predicted values (PV).

We considered a situation in which a given budget (200,000 €) was available to predict the performances of selection candidates with NIRS or genotyping. Depending on the costs of the methods (DNA extraction and genotyping for GS or tissue sampling and NIRS acquisition for PS), we computed the number of selection candidates (*N*) that could be analyzed. The TBV and genomic or NIRS PV of these *N* individuals were then sampled from a multivariate normal distribution with means equal to 0, variances equal to 1 and covariance equal to the square root of reliability (R package mvtnorm (Genz *et al.* 2017)). The expected genetic gain was then computed as the difference between the average TBV of the 400 individuals having the best PV and the average TBV of the population (equal to 0). We selected 400 individuals because for many species, it is feasible to apply heavier phenotyping (multi-environment trials) on a few hundred individuals. We considered two situations; in the first situation, the expected genetic gain of GS and PS was computed for various genotyping and NIRS costs with a reliability set to 0.4. In the second situation, the reliability of GS and PS varied between 0.3 and 0.6, and genotyping and NIRS costs were set to 50 € and 4 €, respectively. For each combination of parameters (reliabilities and costs of GS and PS), the simulation procedure was repeated 1000 times to obtain stable results.

Because genotyping and NIRS costs are highly dependent on the species and the number of samples analyzed, we let the genotyping costs (DNA extraction and genotyping itself) vary between 25 € and 100 € and the NIRS costs (sample treatment and NIRS analysis itself) vary between 1 € and 8 € in the first situation.

To provide concrete examples, we applied this simulation process with the reliabilities and costs that we experienced for wheat and poplar. GS costs were between 35 € and 50 € per individual for wheat and poplar, respectively, and PS costs were between 3 € and 2.5 € per individual for wheat and poplar, respectively. Reliabilities were estimated as the square of predictive abilities estimated by cross-validation divided by the heritability of the adjusted means. For each combination of trait, scenario, and NIRS data considered (tissue, environment), the increase in expected genetic gain using PS instead of GS was computed with the best performing GS model as a reference.

## RESULTS

### Genetic variability captured by NIRS

We first sought to characterize the ability of NIRS to capture genetic variability by estimating genomic heritability and partitioning the variance into genetic (*G*), genetic by environment (*G* × *E*) and residual variances (*e*) along the NIR spectrum collected on a panel of winter wheat (leaves and grains) and a population of black poplar (wood) grown in two contrasting environments. For both species and tissues, genomic heritability was highly variable along the spectrum with peaks above 60%, showing the existence of strong polygenic signals for some wavelengths (Fig. 2, **Fig. S1**). For a given species, the proportion of *G* × *E* variance could reach 24% (poplar), 54% (wheat leaves) or 71% (wheat grains) (Fig. 2). It is interesting to note that for at least half of the wavelengths, the cumulative proportion of *G* and *G* × *E* variances was above 15%, showing that the NIR signal was often partially related to genetics. The kind of tissue analyzed by NIRS seemed to matter, as shown by the comparison of variance partition along spectra obtained on wheat leaves and grains. *G* and *G* × *E* variances were higher and more stable along the spectrum for grains than for leaves.

**Figure 2.**
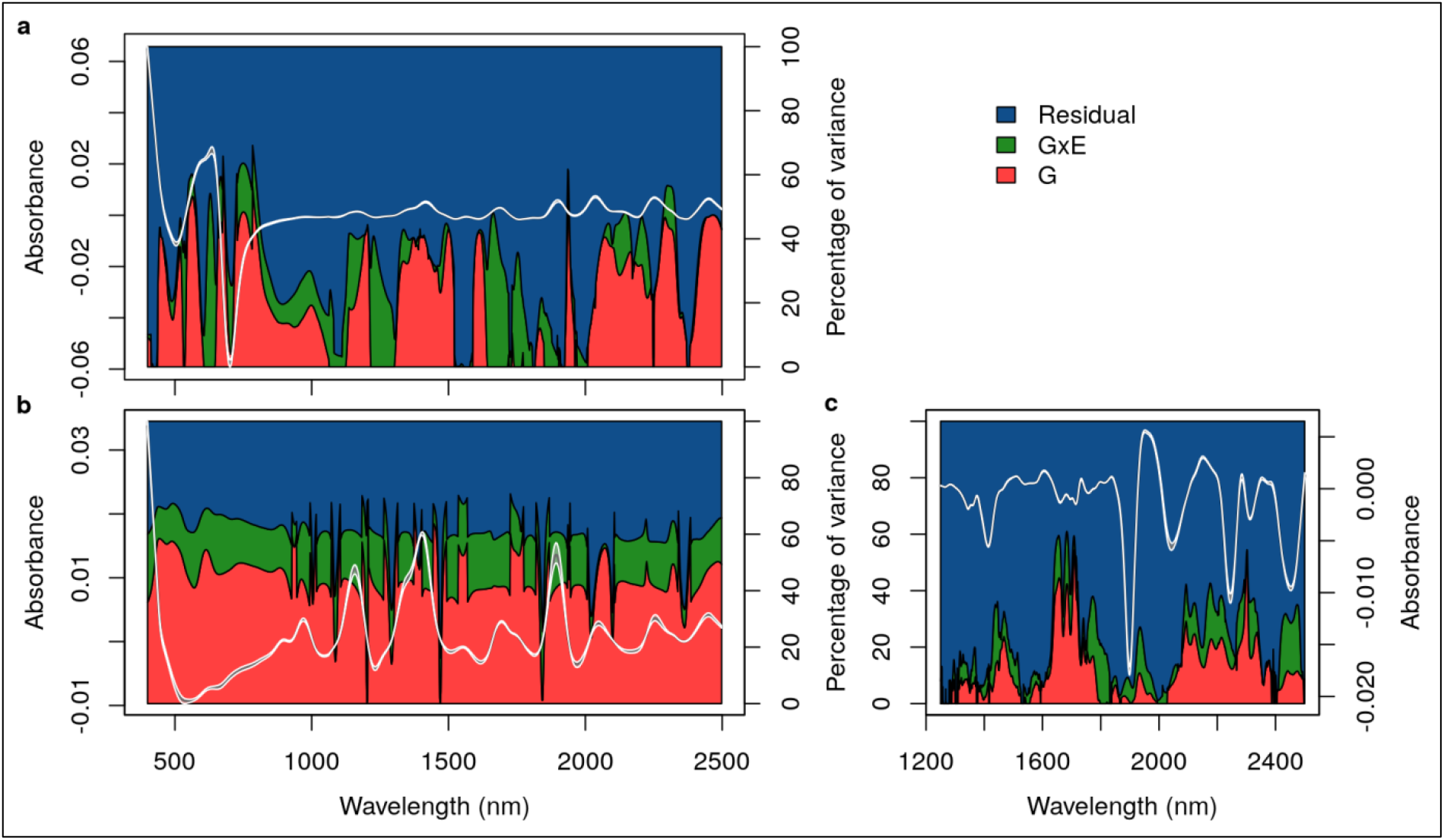
Proportion of genetic (red), genetic by environment (green), and residual (blue) variances along the NIR spectrum of (a) winter wheat leaves, (b) winter wheat grains and (c) poplar wood. NIRS was performed on plant material collected on genotypes grown under favorable and unfavorable environmental conditions. The median normalized and derived spectra, along with their first and third quartiles across the genotypes under study, are indicated in gray.

We ran association mapping along the NIR spectrum to identify wavelengths associated with major QTL (**Fig. S1**). In poplar, the signal appeared to be mainly polygenic with very few QTL detected, and the largest SNP R² was below 0.025 for any wavelength. In contrast, in winter wheat, we detected numerous large-effect QTLs. For some wavelengths, a single SNP could have an R² of 0.23 for leaves and of 0.11 for grains, and this SNP could be in spectrum regions of high or of low genomic heritability (**Fig. S1**). This finding means that depending on the wavelength, NIRS could capture highly polygenic relationships (wavelengths with high genomic heritability) or could tag specific regions of the genome (major QTLs). These two kinds of wavelengths can be useful for making predictions because they can potentially track the two main factors responsible for GS accuracy: relatedness and linkage disequilibrium.

### Comparing predictive abilities obtained with markers and with NIRS

We estimated the efficiency of GS and PS to predict the performance of new individuals within a cross-validation framework. The performances of the individuals in the validation set were predicted with genotypic information in GS (G-BLUP and Bayesian LASSO models) and with NIRS only in PS (RR-BLUP model). We considered two scenarios: in S1, NIRS analysis and cross-validation were performed in the same environment (Fig. 1 a), whereas in S2, the environments in which the cross-validation was applied were different from those in which NIRS was obtained (Fig. 1 b). The broad-sense heritabilities of the adjusted means were above 0.8 for all traits in each environment (**Table S1**).

In wheat, the predictive abilities of PS were highly variable and appeared to be dependent on the predicted trait and on the environment and tissue in which NIRS was measured (Fig. 3 a, b, c, d, Fig. 4). While combining NIRS collected in different environments or different tissues increased the predictive ability, this increase did not occur systematically. One major result is that for both traits, NIRS could lead to better predictions than molecular markers, even in the six independent environments (Fig. 3 c, d, Fig. 4). The gain with NIRS in comparison to molecular markers in S2 was up to 34% and 22% for heading date and grain yield, respectively. In each S2 environment and for both traits, there was always a type of NIRS that performed better or as well as the best GS model (Fig. 4). The gain was even stronger in S1: NIRS led to an increase in predictive ability of up to 53% and 117% for heading date and grain yield, respectively. In poplar, the predictive abilities with NIRS were always lower than those with SNP, except for growth traits under S1 (Fig. 3 e). In the other cases, the predictive ability with NIRS varied depending on the trait and scenario considered, but they were always significantly greater than 0. In general, they were higher when the spectra were collected in the same environment (S1) than when spectra from another environment were used (S2), except for bud flush evaluated in one site and bud set evaluated in another site. Interestingly, irrespectively of the scenario, for some traits apparently unrelated to wood chemical properties, such as resistance to rust or bud set, NIRS predictive abilities were fairly high ranging between 0.34 and 0.53.

**Figure 3.**
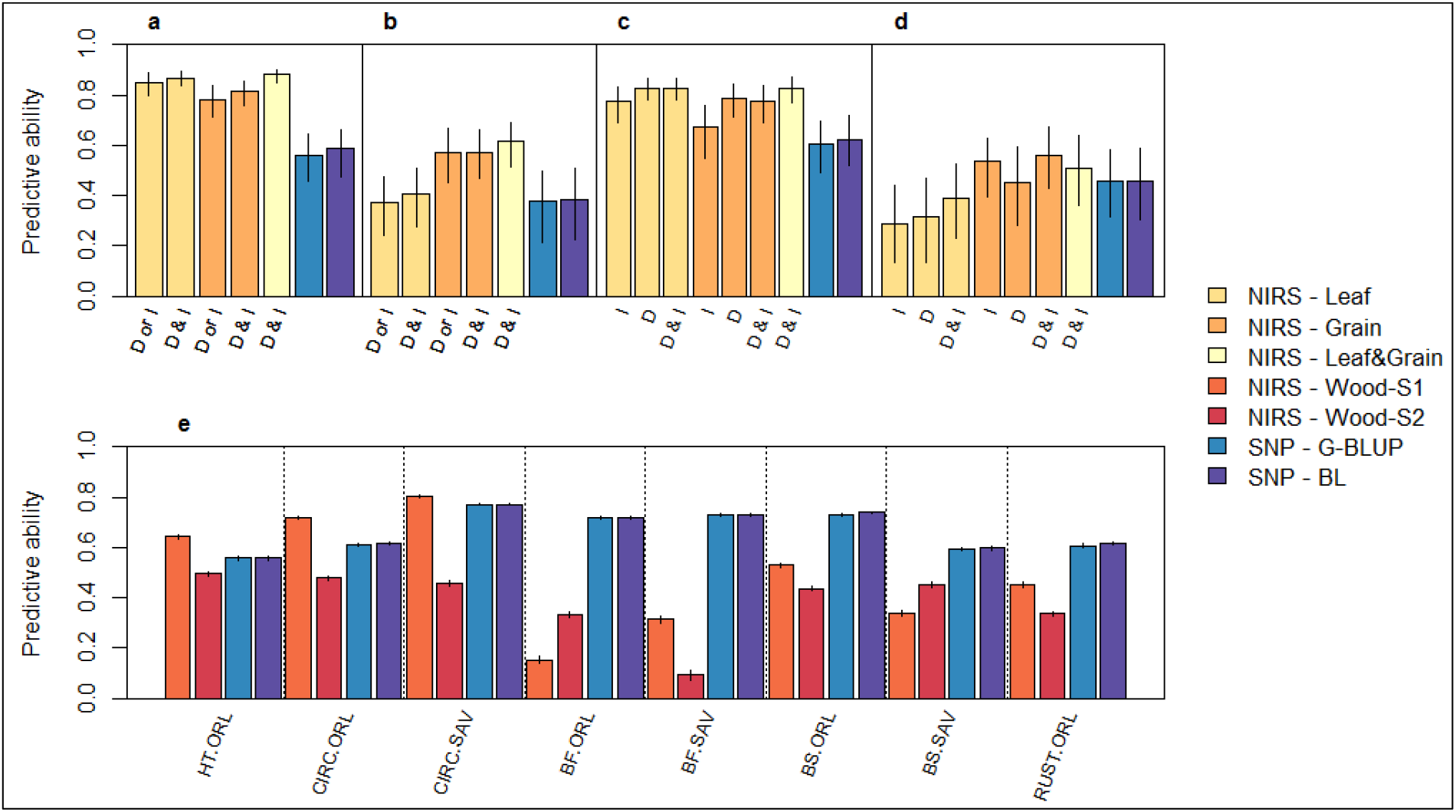
Predictive ability of SNP (G-BLUP or Bayesian LASSO (BL) models) or NIRS (RR-BLUP model) when predicting the phenotypic values of individuals within a cross-validation in winter wheat (a, b, c, d) and black poplar (e). Two scenarios were considered for NIRS prediction: in S1, the RR-BLUP model was trained with NIRS data and phenotypes that were collected within the same environment (a, b, e), whereas in S2, NIRS and phenotypic data used to train the RR-BLUP model were collected in distinct environments (c, d, e). For wheat, two traits were considered: heading date (a, c) and grain yield (b, d). The bars of a, b, c, and d are labeled with the origin of the NIRS data (I: irrigated treatment, D: drought treatment), and the bars of e are labeled with the combination of trait and experiment (HT: height, CIRC: circumference, BF: bud flush, BS: bud set, RUST: resistance to rust, ORL: experimental design in Orléans, France, SAV: experimental design in Savigliano, Italy). The medians of the accuracies obtained over repeated cross-validations are reported as the height of the bars together with the first and third quartiles as confidence intervals.

**Figure 4.**
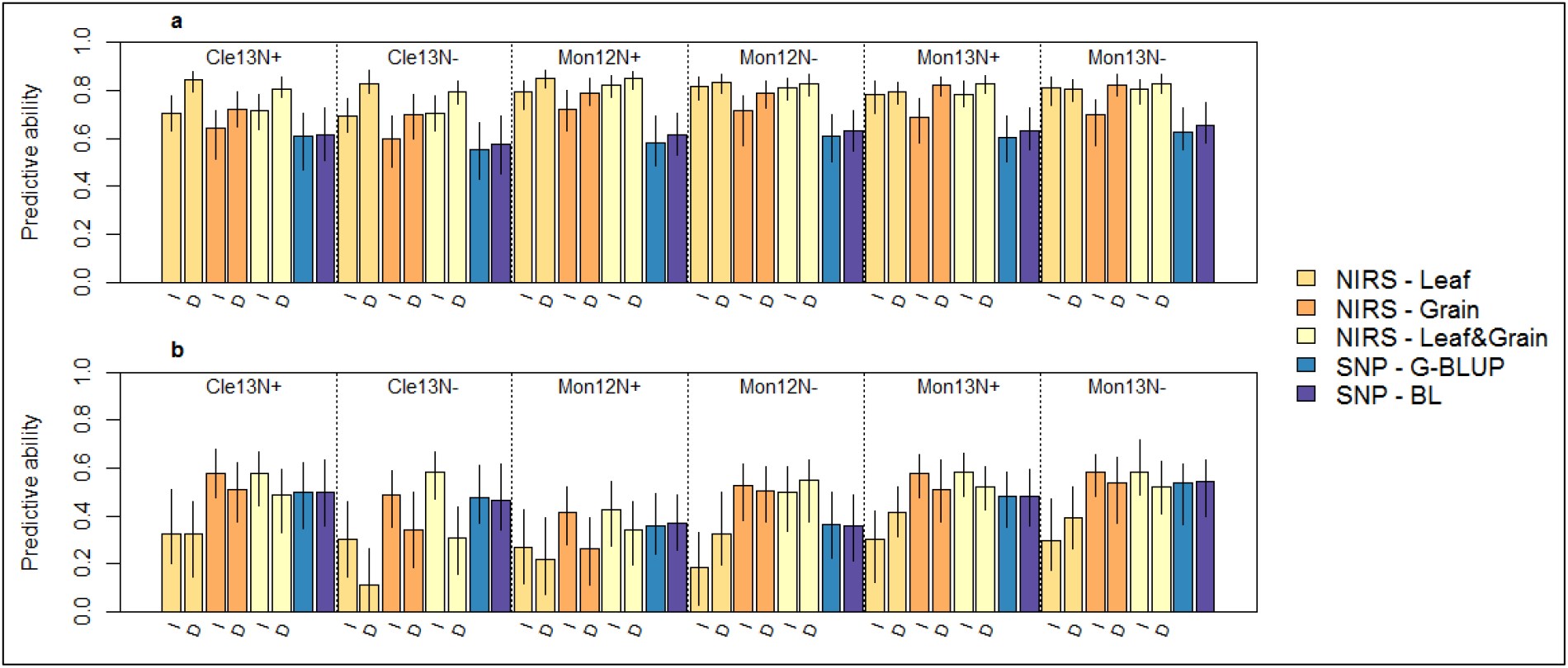
Details of the predictive abilities obtained in scenario S2 for heading date (a) and grain yield (b) for wheat. In S2, the NIRS and phenotypic data used to train the RR-BLUP model were collected in distinct environments. The bars are labeled with the origin of the NIRS data (I: irrigated treatment, D: drought treatment). The medians of the predictive abilities obtained over repeated cross-validations are reported as the height of the bars together with the first and third quartiles as confidence intervals.

### Expected genetic gain with genomic and phenomic selection in a simple example

To further evaluate the potential of PS with respect to GS, the expected genetic gain with both approaches was compared in a simple scenario in which a budget of 200,000 € could be spent to genotype or analyze the NIRS of selection candidates. The difference in efficiency between GS and PS was highly dependent on the genotyping and NIRS costs and on the reliability of the two approaches (Fig. 5). In the scenarios that we considered here, the expected gain of using PS instead of GS was between 11% and 127%. In extreme scenarios in which genotyping was cheap (25 €) and NIRS was expensive (8 €) or in which GS reliability (0.6) was much higher than PS reliability (0.3), PS was still better than GS. We applied the simulation process with the reliabilities and costs obtained in the wheat example (35 € for genotyping and DNA extraction and 3 € for sample treatment and NIRS acquisition). The increase of expected genetic gain with PS in comparison to GS was between + 60% and + 127% for heading date and between −10% and + 222% for grain yield, depending on the tissue and environment used for NIRS acquisition and scenario considered (**Table S2**). In poplar, considering genotyping and NIRS acquisition costs of 50 € and 2.5 €, respectively, as well as the reliabilities estimated with cross-validation predictive abilities, the expected gain in genetic progress varied depending on the trait and scenario considered (**Table S3**). It was mainly positive for growth traits (−2 to 93%), bud set (−6 to 25%) and rust resistance (−10 to 21%), whereas for bud flush, NIRS prediction did not seem to provide any advantage over regular SNP-based prediction.

**Figure 5.**
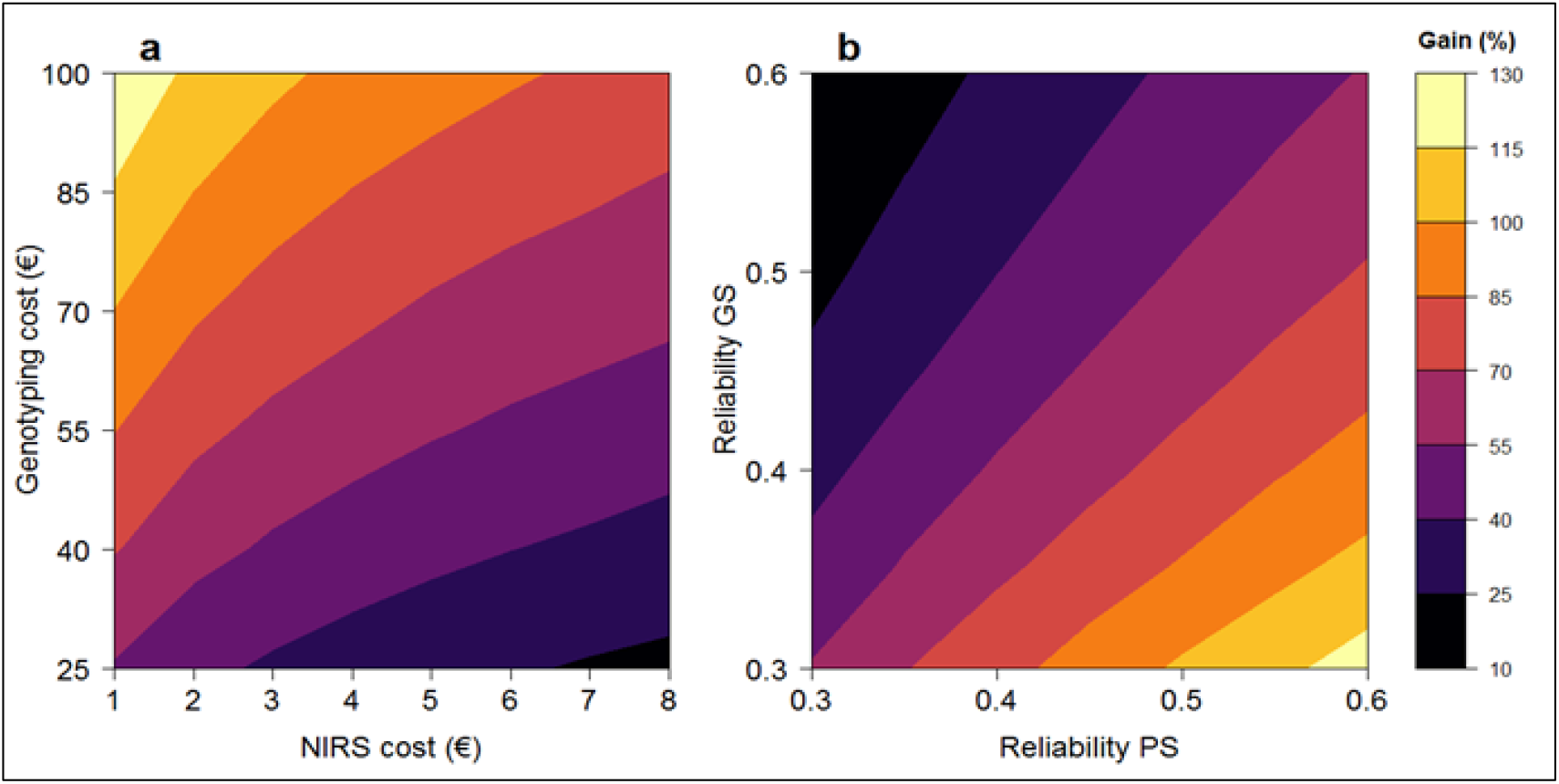
Theoretical increase of expected genetic gain (%) by using NIRS instead of genotyping. a: Expected genetic gain for various genotyping and NIRS costs for a reliability of 0.4, a budget of 200,000 €, and a selection of 400 individuals. b: Expected genetic gain for various reliabilities, a budget of 200,000 €, genotyping and NIRS costs of 50 € and 4 €, respectively, and a selection of 400 individuals. For each scenario, true breeding values and estimated breeding values were simulated thanks to multivariate normal distributions with a covariance adapted to the chosen reliability.

## DISCUSSION

In typical plant breeding programs, breeders have to select among thousands to millions of individuals. For most individuals, this selection is often based on a very small amount of phenotypic information because it is too expensive or simply impossible to make a precise phenotypic evaluation. It is also difficult and too expensive to genotype all individuals to apply GS, despite important economies of scales. Alternative approaches based on endophenotypes such as transcriptomes or metabolomes have been proposed to predict phenotypes (Fu *et al.* 2012; Riedelsheimer *et al.* 2012; Feher *et al.* 2014; Ward *et al.* 2015; Fernandez *et al.* 2016; Guo *et al.* 2016; Xu *et al.* 2016; Zenke-Philippi *et al.* 2016; Westhues *et al.* 2017; Seifert et al. 2018; Schrag *et al.* 2018), but their relatively low throughput and high costs are still likely to hamper their deployment at a large scale. To increase genetic progress in this context, we propose a new approach in which we use NIRS as high-throughput phenotypes to make predictions at low costs. The basic idea of this approach, which we call “phenomic selection” (PS), is that the absorbance of a sample in the near-infrared range is mainly related to its chemical composition, which depends itself on endophenotypes and genetics. Therefore, NIRS is supposed to capture at least part of the genetic variance, and as a result, one could use it to make predictions of traits unrelated to the analyzed tissue or in independent environments. The process of PS is similar to GS, but instead of reference material and selection candidates being genotyped, they are analyzed by NIRS.

We applied PS to the NIR spectrum of different tissues sampled on an association population of poplar and a panel of elite winter wheat. By estimating the extent of genetic variance along the NIR spectrum of poplar wood and winter wheat leaves and grains, we could show that most wavelengths captured part of the genetic variability (Fig. 1). This result agrees with previous findings with eucalyptus wood (Hein and Chaix 2013), but whether this will still be true within pedigrees with a narrower genetic basis remains to be assessed. O’Reilly-Wapstra *et al.* (2013) have shown that NIR spectra collected on eucalyptus leaves could differentiate full-sibs, even though the extent of genetic variation captured was lower than at the inter-specific level. Still these results suggest that NIRS could be valuable to capture some Mendelian sampling and that PS would work within pedigrees, but this hypothesis should clearly be tested in future work.

In the present work, the NIR spectra were specific to the environments in which they were obtained, but when they were analyzed jointly, we observed that *G* variance was superior to *G* × *E* variance for most wavelengths in both species. Posada *et al.* (2008) also reported a similar trend with coffee grains. This finding shows that even if the absorbances were partly environment specific, it should be possible to make predictions in independent environments. This result was further demonstrated by the good predictive abilities obtained with PS for most phenotypes in both species in scenario S2, *i.e.*, when the environment in which we trained the calibration model was different from the environment in which we collected NIRS. For both species, PS abilities were in the same range as GS abilities, sometimes performing better and sometimes performing worse than one another. For wheat, the results were very encouraging as we always found a situation (combination of environment and tissue analyzed) for which NIRS performed better than GS, even in six independent environments. More importantly, even when the correlation between the S1 and S2 environment was as low as 0.16 for the predicted trait (**Table S4**, GY in Mon12N-), PS could produce better predictions than GS (Fig. 4). In other words, a relationship matrix computed with NIRS obtained in one specific environment could be used to make predictions in completely different environments. These promising results obtained in scenarios S1 and S2 open the way to important opportunities in the plant breeding community. As revealed by our theoretical computations (Fig. 5), we expect PS to be able to generate large gains in genetic progress in comparison to GS, even in pessimistic scenarios. In the realistic scenarios that we experienced, the expected gain brought by using PS instead of GS could be up to 81% for wheat grain yield in scenario S2 (**Table S2**).

Nevertheless, these simulations have to be considered with caution, because of the strength of the underlying hypotheses. Our work has shown the interest of the proposed PS approach within a given generation that may clearly be applicable within plant breeding programs to assess the performance of the candidate for selection. But, more work is clearly needed to establish the proportion of the variance captured by NIRS (and by endophenotypes) that is heritable in the narrow-sense and thus transmitted to the next generations to be further used at different stages of the breeding programs or in different breeding contexts. Indeed, similarly to endophenotypes we expect NIRS to capture non-additive genetic effects which may overestimate the expected genetic progress across multiple generations. Nevertheless on the other side it can be highly valuable to predict phenotypes, including the effect of interactions and regulatory networks, at key steps of the breeding schemes. In plant breeding, one major objective during the first generations is to produce numerous individuals with the same genotypes (by autofecondation, doubled haploid techniques or clonal reproduction) to allow for field evaluation in multi-environment trial (MET). Because this field evaluation is the most expensive step in the breeding schemes and because it is applied on replicable genotypes, PS would be of major interest to select among all candidates the genotypes that will be evaluated in the MET. In this situation, predicting phenotypes instead of additive values is clearly an advantage as the same genotypes can be replicated in many individuals. Another important question related to the efficiency of PS across multiple generations is about the frequency of formula update to maintain a sufficient level of accuracy. This question, which is also relevant for GS, must be addressed in future work. We also believe that future work should assess the efficiency of PS with NIRS obtained from tissues collected on young plants. Ideally, we would like PS to be efficient with NIRS collected on the youngest possible plant to have the information as early as possible and at low cost. We could show for wheat that for fixed material, NIRS collected on seeds, so before sowing, was efficient to run PS, which offers very interesting perspectives for this species. The studies on endophenotypic variations in maize (Riedelsheimer *et al.* 2012; Fu *et al.* 2012; Guo *et al.* 2016; Schrag *et al.* 2018), rice (Xu *et al.* 2016) and wheat (Ward *et al.* 2015) also demonstrated that the characterization of germinated seeds or seedlings was efficient to estimate kinships resulting in accurate predictions. These results are promising, but this needs to be tested for other species and on other datasets.

There are various applications of PS, which we see both as a complement and as an alternative to GS depending on the situation. The first obvious application of PS is its use when no genotyping tool is available at a reasonable cost, which is still the case for many orphan organisms. For these species, PS could potentially be a new efficient breeding tool to increase genetic progress. As mentioned before, a second application would be to use PS to screen nearly fixed material or clones, as PS (in the same manner as selection on endophenotypes) is likely to capture non-additive genetic effects. Even if the prediction accuracy is low, PS can be used to filter out a given proportion of selection candidates. One should define this proportion with respect to PS accuracy: the higher the accuracy, the more confident we are at filtering out many individuals without losing the best candidates. Note that even if PS is less accurate than GS, it could nevertheless be interesting to filter out the worst individuals considering the low cost of NIRS acquisition, and the fact that NIRS is often already routinely carried out (for example, in cereals or forest trees to predict quality traits). In a second step, one could use GS to make complementary predictions on a limited number of selection candidates. A last major application of PS would be to help conservation geneticists manage diversity collections. The use of genotyping to organize seed banks and to screen and define core collections is strongly limited by its cost. PS offers a new opportunity to manage seed banks because it allows distance matrices to be computed cheaply and reliably.

Considering that PS gave interesting results for both a tree and an annual crop regarding various traits related to development, productivity and tolerance to disease and using tissues of a completely different nature (wood, leaf, grain), we can expect PS to work in many other plants and possibly in animal species using NIRS on organic tissues or fluids. Our work constitutes a proof of concept and a first attempt at PS, which clearly opens new perspectives for the breeding community. Indeed, one could further optimize many parameters to increase PS efficiency. The differences observed here between the PS efficiencies reported for wheat and poplar could represent a first direction for improving the approach. Indeed, PS appeared to be more efficient in wheat than in poplar and several hypotheses could be proposed to explain this result. First, spectra were acquired on different spectrometers resulting in a broader wavelength range in wheat, which also covered the visible part of the electromagnetic spectrum. Consequently, the information brought by the spectra on wheat tissues was potentially richer than the one brought by the spectra on poplar. Second, we could see that in wheat a larger proportion of *G* and *G* × *E* variance could be captured by the spectra regardless of the tissue sampled and that this was especially true for the lowest wavelengths (including the visible part), which were absent in poplar. Third, the tissues in which NIRS was collected differed, and this difference seems to be an important parameter as highlighted by the differences in predictive ability between leaf and grain in wheat.

Another possibility for the improvement of PS efficiency could be the optimization of the growing conditions of plants in the reference experiment. In wheat, it was typically better to use NIRS collected on plants grown in unfavorable conditions than in favorable conditions. This result might be explained by more pronounced dissimilarities between genetically distant individuals in conditions of stress. Therefore, there is a clear need to optimize these conditions. Once the NIRS data are collected, one could also try to improve the pretreatment of the signal and the statistical model of calibration. In our case, we choose as pretreatment the first derivative of the normalized spectrum, but other options could be tested, and these options might not necessarily be the same depending on the species considered, environment, tissue sampled or target trait. For calibrations, we have used RR-BLUP, but one might test other techniques, such as those typically allowing non-additive effects or involving feature selection, to improve the accuracy of PS. These points clearly indicate that there is great room of improvement of PS, which will likely constitute in the near future an active field of research. Finally, the recent advent of portable NIR devices as well as of hyperspectral imaging allows this technology to be used in the field. Unmanned vehicles and robots are currently being developed and can already be used to automatically collect reflectance at an industrial scale (Madec *et al.* 2017; Aguate *et al.* 2017). These new developments will considerably increase the throughput and conversely decrease the cost of NIRS data. We thus expect that these technological advances will reinforce the advantages of the proposed PS.

## Data availability

The datasets generated during and/or analysed during the current study are available in the INRA Dataverse repository (https://data.inra.fr/). They can be accessed with the following link: http://dx.doi.org/10.15454/MB4G3T.

## Code availability

R code used throughout the study is available upon request.

## ACKNOWLEDGEMENTS

The authors gratefully acknowledge the staff of the INRA GBFOR experimental unit for the establishment and management of the poplar experimental design in Orléans, the collection of wood samples in each site, and their contribution to phenotypic measurements on poplars in Orléans; Alasia Franco Vivai staff for management of the poplar experimental plantation in Savigliano, and M. Sabatti and F. Fabbrini for their contribution to phenotypic measurements on poplars in Savigliano. We acknowledge the staff of the INRA GenoBois platform for the preparation of samples and collection of NIRS on wood samples and the staff of EPGV and BioForA for their contribution to obtaining SNP data on poplar. We would like to thank J. Messaoud for NIRS acquisition on wheat samples, V. Allard, B. Adam and D. Cormier for implementation of the rain-out shelter experiment (Phéno3C, INRA Clermont-Ferrand), and E. Heumez (UE GCIE) for the experiment in Estrées-Mons. We would also like to thank A. Chateigner, L. Sanchez, G. Charmet, V. Allard, P. Martre, L. Inchboard and S. Bouchet for useful discussions and comments on the manuscript. We are grateful to the Genotoul bioinformatics platform Toulouse Midi-Pyrenees (Bioinfo Genotoul) for providing computing resources. Establishment and management of the poplar experimental sites until harvests were carried out with financial support from the NOVELTREE project (EU-FP7-211868). NIRS measurements on poplar wood samples were supported by the SYBIOPOP project funded by the French National Research Agency (ANR-13-JSV6-0001). Management of the wheat multi-environment trials was financially supported by the French National Research National Agency under Investment for the Future (BreedWheat project ANR-10-BTBR-03) and by FranceAgriMer. The Phéno3C platform was financially funded by the French National Research National Agency under the Investment for the Future Phenome project (ANR-11-INBS-12) and by the European Regional Development Fund (AV0011535).

## AUTHOR CONTRIBUTIONS

V.S. and R.R. designed the study, analyzed the data and wrote the paper with input from J-P.C., P.F.R, E.P., J.L.G., and C.B.

## SUPPLEMENTARY DATA

### Supplementary figures

**Figure S1.**
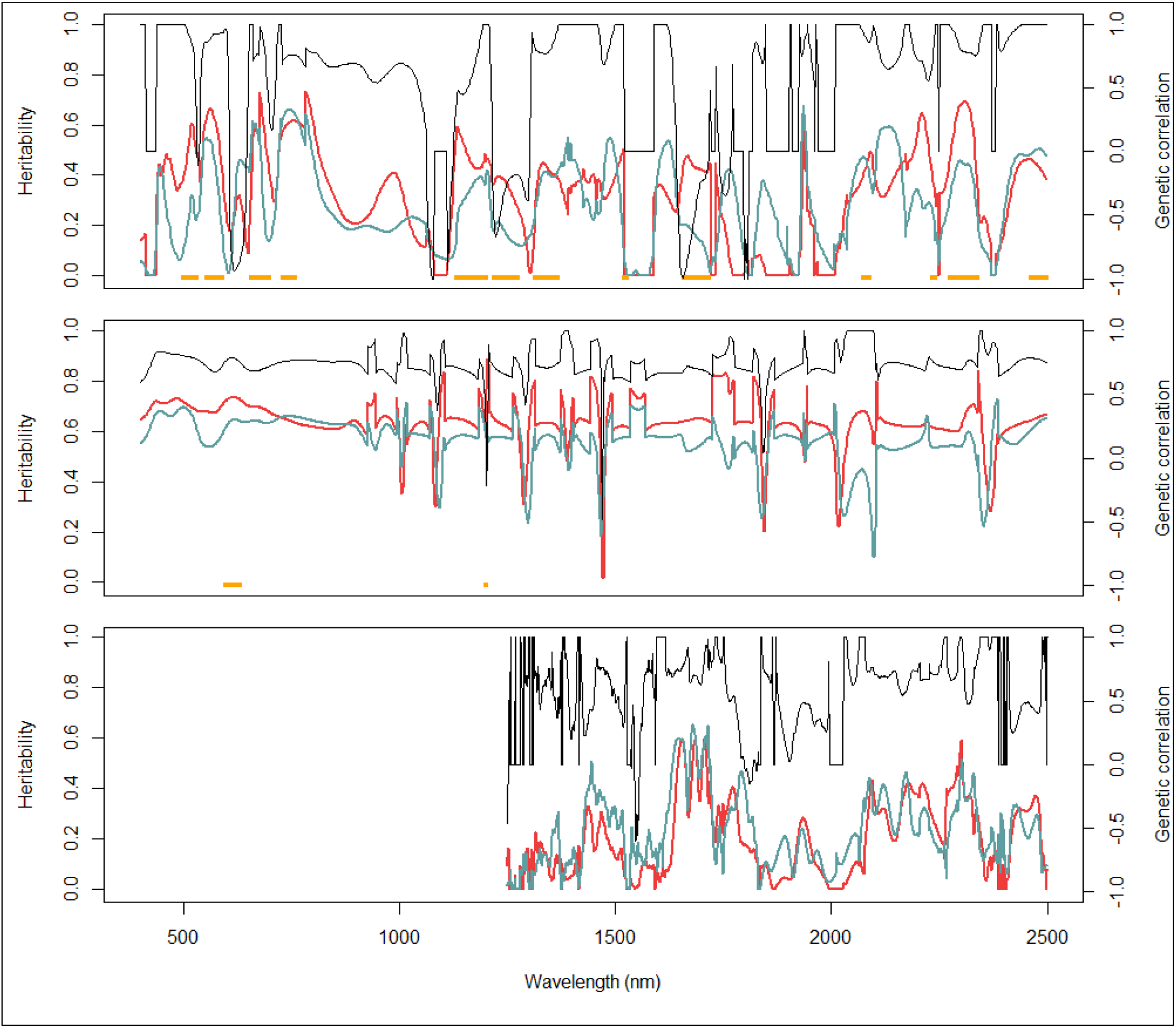
Genomic heritability (color) and genetic correlation (black) along spectra collected on winter wheat leaves (a), winter wheat grains (b) and poplar wood (c). The genotypes were grown under two environmental conditions, unfavorable (red) and favorable (blue). The wavelengths at which absorbance is associated with at least one SNP having a major effect (R² higher or equal to 10%) are indicated with orange dots at the bottom of each graph.

### Supplementary tables

**Table S1:** Broad-sense heritabilities of the adjusted means.

| Species | Location | Year | Treatment | Code environment | Trait identifier | Trait name | Heritability |
| --- | --- | --- | --- | --- | --- | --- | --- |
| Wheat | Clermont-Ferrand | 2016 | Irrigated | IRR | HD | Heading date | 0.94 |
| Wheat | Clermont-Ferrand | 2016 | Irrigated | IRR | GY | Grain yield | 0.78 |
| Wheat | Clermont-Ferrand | 2016 | Drought | DRY | HD | Heading date | 0.94 |
| Wheat | Clermont-Ferrand | 2016 | Drought | DRY | GY | Grain yield | 0.75 |
| Wheat | Estrée-Mons | 2012 | N+ | Mon12N+ | HD | Heading date | 0.97 |
| Wheat | Estrée-Mons | 2012 | N+ | Mon12N+ | GY | Grain yield | 0.81 |
| Wheat | Estrée-Mons | 2012 | N- | Mon12N- | HD | Heading date | 0.96 |
| Wheat | Estrée-Mons | 2012 | N- | Mon12N- | GY | Grain yield | 0.83 |
| Wheat | Estrée-Mons | 2013 | N+ | Mon13N+ | HD | Heading date | 0.98 |
| Wheat | Estrée-Mons | 2013 | N+ | Mon13N+ | GY | Grain yield | 0.85 |
| Wheat | Estrée-Mons | 2013 | N- | Mon13N- | HD | Heading date | 0.99 |
| Wheat | Estrée-Mons | 2013 | N- | Mon13N- | GY | Grain yield | 0.86 |
| Wheat | Clermont-Ferrand | 2013 | N+ | Cle13N+ | HD | Heading date | 0.98 |
| Wheat | Clermont-Ferrand | 2013 | N+ | Cle13N+ | GY | Grain yield | 0.86 |
| Wheat | Clermont-Ferrand | 2013 | N- | Cle13N- | HD | Heading date | 0.90 |
| Wheat | Clermont-Ferrand | 2013 | N- | Cle13N- | GY | Grain yield | 0.88 |
| Poplar | Orléans | 2011 | - | ORL | HT | Height | 0.88 |
| Poplar | Orléans | 2011 | - | ORL | CIRC | Circumference | 0.86 |
| Poplar | Orléans | 2009 | - | ORL | BF | Bud flush | 0.96 |
| Poplar | Orléans | 2009 | - | ORL | BS | Bud set | 0.97 |
| Poplar | Orléans | 2009 | - | ORL | RUST | Rust resistance | 0.89 |
| Poplar | Savigliano | 2010 | - | SAV | CIRC | Circumference | 0.92 |
| Poplar | Savigliano | 2010 | - | SAV | BF | Bud flush | 0.90 |
| Poplar | Savigliano | 2010 | - | SAV | BS | Bud set | 0.92 |

**Table S2:** Increase of expected genetic gain (%) by using PS instead of GS for wheat. The expected genetic gain of PS and GS was estimated with the estimated heritabilities, the costs that we experienced (3 € and 35 € for PS and GS, respectively) and the predictive abilities obtained in cross-validation in scenarios S1 and S2. For each combination of scenario, trait, and NIRS data considered (tissue and environment), the increase of expected genetic gain of PS was estimated with the best performing GS model as a reference.

|  | S1 |  |  |  | S2 |  |
| --- | --- | --- | --- | --- | --- | --- |
|  | GY-IRR | GY-DRY | HD-IRR | HD-DRY | GY | HD |
| max | 94 | 222 | 121 | 127 | 81 | 98 |
| min | -2 | 113 | 80 | 106 | -10 | 60 |

**Table S3:**
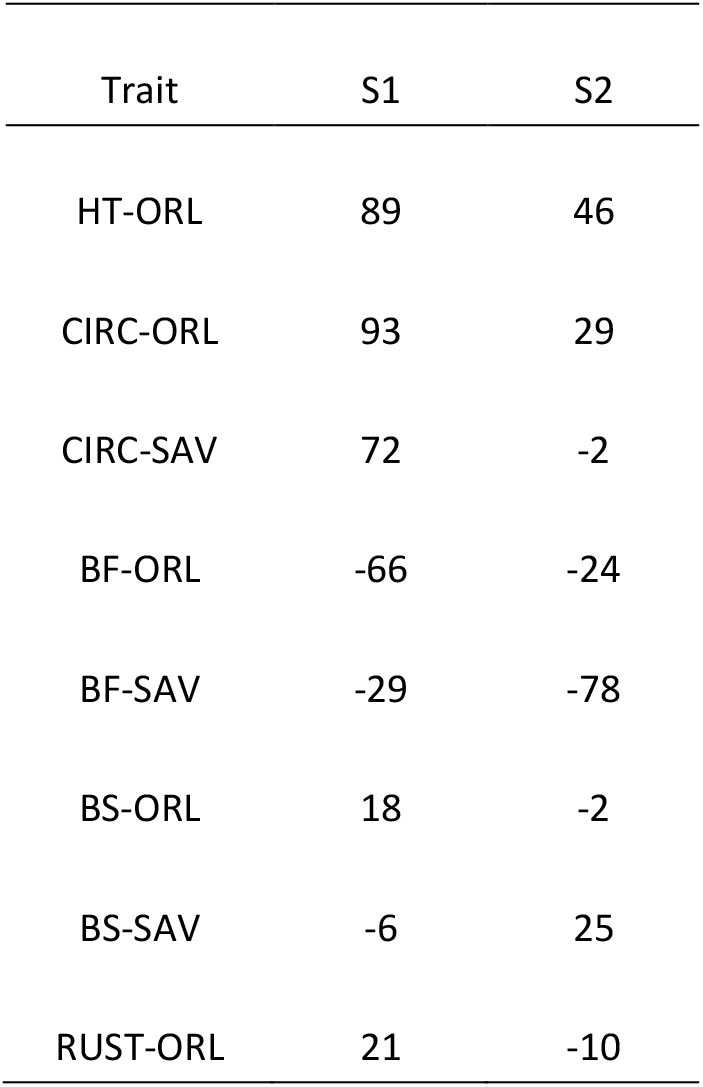
Increase of expected genetic gain (%) by using PS instead of GS for poplar. The expected genetic gain of PS and GS was estimated with the estimated heritabilities, the costs that we experienced (2.5 € and 50 € for PS and GS, respectively) and the predictive abilities obtained in cross-validation in scenarios S1 and S2. For each combination of scenario and trait, the increase of expected genetic gain of PS was estimated with the best performing GS model as a reference.

**Table S4:**
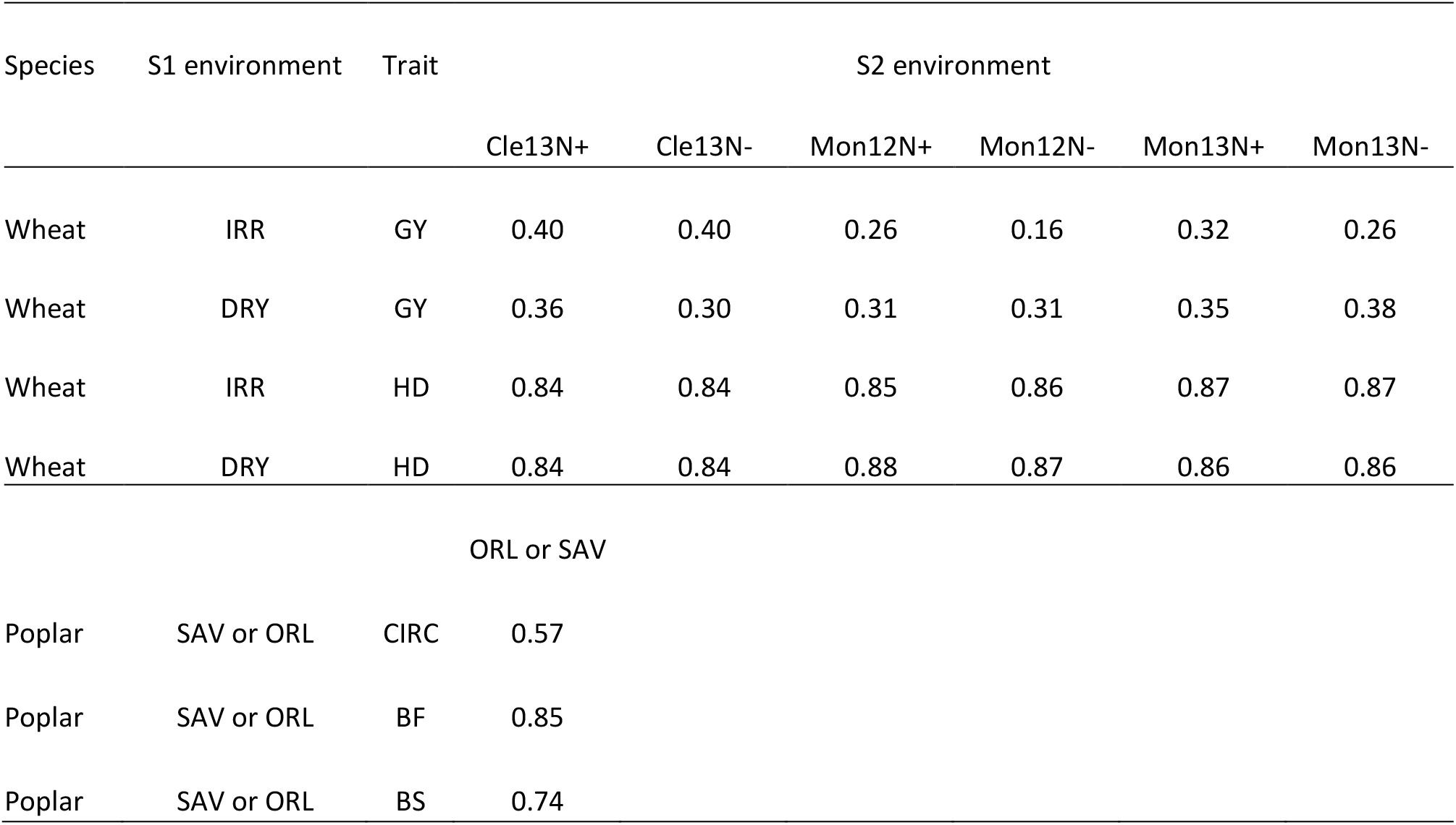
Correlations of traits between S1 and S2 environments.

